# Increased disease burden in Interleukin-3 deficient mice after *Mycobacterium tuberculosis* and herpes simplex virus infections

**DOI:** 10.1101/2021.03.07.434271

**Authors:** Shajo Kunnath-Velayudhan, Tony W. Ng, Neeraj K. Saini, Michael F. Goldberg, Pooja Arora, Jiayong Xu, John Kim, Betsy C. Herold, John Chan, William R. Jacobs, Steven A. Porcelli

## Abstract

Interleukin-3 (IL-3) is produced during infections caused by parasites, bacteria and viruses, but its contribution to immunity in this context remains largely unknown. In mouse models of parasitic infections, in which the effects of IL-3 have been most extensively studied, IL-3 has been variously reported as protective, detrimental or inconsequential. Similarly, mixed results have been reported in viral and bacterial infection models. Here, we investigated the effects of IL-3 in mouse models of *Mycobacterium tuberculosis* and herpes simplex virus type 1 (HSV-1) and type 2 (HSV-2) infections by assessing the pathogen burden, disease manifestations and survival following infection. After infection with *M. tuberculosis,* IL-3 deficient mice showed higher bacillary burden, increased lung pathology and reduced survival compared to wild type mice. After infection with HSV-1 through cutaneous route and HSV-2 through vaginal route, IL-3 deficient mice showed higher viral burden, increased disease manifestations and reduced survival compared to wild type mice. Our results show that IL-3 makes a subtle but significant contribution to protective immunity in these mouse models of bacterial and viral infections.

## 1 Introduction

The function of interleukin-3 (IL-3), a cytokine produced by many hematopoietic cells including activated T cells and non-hematopoietic cells, remains mostly unknown (1–3). IL-3 deficient mice display normal steady-state hematopoiesis, despite the fact that IL-3 promotes growth and proliferation of hematopoetic progenitor cells (4). The role of IL-3 in infectious diseases has been investigated using IL-3 deficient mice or mice receiving anti-IL-3 neutralizing antibodies. These experiments were performed mostly in parasitic infection models given its effect on proliferation of mast cells and basophils, and in potentiation of Th2 immunity (5). However, these studies yielded contradictory results on whether IL-3 is protective or deleterious in the context of parasitic infections. For example, expulsion of *Stronglyoides venezuelensis* was impaired in IL-3-deficient mice compared to wild type mice suggesting a protective role of IL-3 (5). In contrast, lower levels of *Plasmodium berghei* and prolonged survival was seen in IL-3 deficient male but not female mice compared to wild type mice, suggesting a deleterious role of IL-3 in this context (6). The deleterious role of IL-3 was also indicated by earlier studies on cutaneous leishmaniasis, and in combination with GM-CSF neutralization in *Plasmodium berghei* infection (7–9). Studies on *Nippostrongylus brasiliensis* on the other hand did not show any protective or deleterious role of IL-3, suggesting that its effect varies for different parasites (5). Similar conflicting results have been obtained in non-parasitic infection models, such as experimental infections with herpes simplex virus type 1 (HSV-1) or *Listeria monocytogenes* in which IL-3 has been protective or inconsequential, respectively (10, 11). Thus, in contrast to non-infection models such as experimental autoimmune encephalitis and myocarditis, lupus nephritis, delayed type hypersensitivity and sepsis where the effect of IL-3 has been consistently deleterious (1, 4, 12–14), the role of IL-3 in infections is less uniform and may vary depending on the particular microorganisms involved.

We recently reported that in mice, infections with various species of mycobacteria, *Listeria monocytogenes* and HSV-2 induce IL-3 secreting T helper cells (15, 16). These cells were generated by infection at the skin and mucosa but not by infections introduced directly into the blood. The pattern of co-expression of other cytokines by these cells was also distinct, with most IL-3 producing T cells co-secreted GM-CSF. These cytokines share the beta subunit of their respective receptors, and their transcription is controlled by similar regulatory sequences (17). IL-3-secreting T cells also co-secreted IFNγ, the signature cytokine of Th1 helper cells, and additional cytokines that define multifunctionality including IL-2 and TNF (15). Generation of IL-3 producing T cells *in vitro* showed that their differentiation from naïve CD4^+^ T cells was dependent on IL-1 family cytokines, and was inhibited by cytokines that induce canonical Th1 or Th2 cells (15). The characteristic cytokine expression pattern of these cells, their dependence on initial stimulation by antigens introduced at the barrier surfaces, and the unique cytokine milieu driving their generation suggested that IL-3 secreting CD4^+^ T cells are a distinct and functionally specialized subset of T helper cells. Since IL-3 secreting T helper cells show overlap in their cytokine secretion profile with conventional Th1 helper cells which are known to be important in protection against mycobacterial and herpetic infections (18, 19), we hypothesized that IL-3 produced by these cells could have a protective function in these infections. Our results in the current study showed that IL-3 has a protective role in infections caused by *M. tuberculosis* and HSV in mice.

## 2 Methods

### Mice

Six- to eight-week-old female wild-type (WT) C57BL/6 and BALB/c mice were purchased from The Jackson Laboratory. Mice with homozygous targeted deletion of the *IL-3* gene on the BALB/c background (BALB/c *IL-3* ^−/−^) were obtained from Dr. Booki Min (Cleveland Clinic, OH) (4) and maintained by breeding in our facility. These mice were backcrossed for 10 generations to obtain *IL-3* ^−/−^ mice in the C57BL/6 background (C57BL/6 *IL-3* ^−/−^). Either BALB/c *IL-3* ^−/−^ or C57BL/6 *IL-3* ^−/−^ mice were used in experiments as indicated in the description of individual experiments. All mice were maintained in specific pathogen-free conditions. All procedures involving the use of animals were in compliance with the protocols approved by the Einstein Institutional Animal Use and Biosafety Committee.

### Antibodies and cell lines

Neutralizing antibody to IL-3 was generated in our laboratory from the MP2-8F8 hybridoma cell line obtained from Dr. Fred D. Finkelman (University of Cincinnati, OH) (20) and purified from culture filtrates by standard protein G column chromatography (Amersham Biosciences/GE Healthcare, Piscataway, NJ). The hybridoma cells were grown in RPMI-1640 medium (Life Technologies) supplemented with penicillin-streptomycin (Life Technologies), immunoglobulin-depleted FBS (10%; Atlanta Biologicals), β-mercaptoethanol (Life Technologies), and essential and nonessential amino acids (Life Technologies). The IL-3 dependent cell line M-NFS-60 was obtained from ATCC and was used to assess neutralization by anti-IL-3 antibody (20). M-NFS-60 cells were grown in DMEM medium (Life Technologies) with supplements as describe above. MTT assay was performed as previously described (21). Matched isotype control antibodies were purchased from Bio X Cell (Lebanon, NH). Wild type C57BL/6 mice were used for all IL-3 neutralization experiments. For this, each mouse was injected intraperitoneally with 400 μg of anti-IL-3 antibody (clone MP2-8F8) or matched isotype control antibody twice per week till sacrifice starting from the day prior to infection, or in vaccinated mice, starting from the day prior to vaccination.

### Infection with *Mycobacterium tuberculosis*

*M. bovis* BCG-Danish (Statens Serum Institute, Copenhagen, Denmark) and *M. tuberculosis* (strain H37Rv) were used for vaccination and challenge, respectively. All experiments involving *M. tuberculosis* were conducted in biosafety level 3 conditions. Details of bacterial culture and infection have been described previously (15). Briefly, bacteria were harvested from liquid cultures and used to prepare frozen stocks that were subsequently used to make single cell suspensions immediately before vaccination or challenge. Mice were vaccinated with 2×10^6 CFU of *M. bovis* BCG-Danish subcutaneously at the base of the tail. Four weeks after vaccination, mice were challenged through the aerosol route with *M. tuberculosis* (strain H37Rv) as previously described (15). Unless otherwise specified, approximately 100 bacteria were deposited into the lungs of each animal as confirmed by plating of whole lung homogenates on Middlebrook 7H10 agar at 24 hours post-aerosol exposure. At four weeks post-challenge, unless otherwise specified, mice were euthanized, and organs were harvested for bacterial enumeration. For survival experiments, BAL/c *IL-3* ^−/−^ and control mice were challenged with *M. tuberculosis* by aerosol route as described above or intravenously (1 x 10^7 CFU) and monitored until death or sacrifice point.

For histopathological examination, the organs were harvested and fixed in 10% neutral buffered formalin (Fisher Scientific, Fair Lawn, NJ). Tissues were embedded with paraffin, sectioned at 5 μm thickness, and stained with hematoxylin and eosin. The area of granulomatous involvement was assessed by scanning 20 random fields in 3 sections per mouse. The lymphocytic infiltrates in the granulomas were graded visually by a trained veterinary pathologist as follows: 0 = none; 1 = minimal; 2 = mild; 3 = moderate; 4 = marked; 5 = severe.

### Infection with HSV

Vaginal infections were performed with a clinical isolate of HSV-2 (strain 4674) and skin infections were performed with HSV-1 (strain B^3^ × 1.1) as described previously (15, 22, 23) Briefly, the viruses were propagated on Vero cells (CCL-81; ATCC), and viral stocks with known titer were prepared and frozen, and subsequently thawed and diluted in PBS on the day of infection. Each mouse was infected with a dose of virus previously determined to cause death in 90% of the animals (LD90; 5 x 10^4 PFU for HSV-2 intra-vaginal infections and 1 x 10^5 PFU for HSV-1 skin infections). For intravaginal infections, mice were injected with 2.5 mg of medroxyprogesterone acetate (Sicor Pharmaceuticals) s.c. at the scruff of the neck 5 days prior to the infection. After infection, the mice were monitored for 14 days for signs of disease, and viral titers were quantified from vaginal swab samples as described previously (24). Epithelial disease for intravaginal infections was scored as follows: (1) mild erythema, (2) hair loss, erythema, edema, (3) severe edema, hair loss, lesion formation, (4) severe ulcerations, multiple lesions and (5) death. Neurological disease for intravaginal infection was scored as follows: (1) urinary retention, (2) urinary retention and constipation, hind-limp paresis, (3) hind-limb paralysis (one leg), (4) complete hind limb paralysis (both legs), and (5) death. Mice were euthanized at a score of 3 or 4 and assigned a score of 5 on subsequent days for analyses. Severity of disease in the skin scarification model was scored as follows: (1) primary lesion or erythema at site of inoculation; (2) distant site zosteriform lesions with mild edema or erythema, (3) severe ulceration and edema with increased epidermal spread, (4) hind-limb paralysis and (5) death. Mice that were euthanized at a score of 4 were given a value of 5 the next day.

### Statistical analysis

For pairwise comparisons, Student’s unpaired t tests were performed to test for significance and reported as significant for p < 0.05. For multiple comparisons, a one-way ANOVA was performed initially to test for significance of overall difference among group means. If ANOVA resulted in p values < 0.05, post hoc comparisons were performed as Student’s unpaired t tests and indicated in the figure legends. Log-rank tests were used to compare survival curves. All statistical analyses were performed using GraphPad Prism software, version 8.

## 3 Results

### 3.1 Role of IL-3 against infection with *M. tuberculosis*

To examine the role of IL-3 in a mouse model of tuberculosis, we first used IL-3 blocking antibodies to neutralize IL-3 function during *M. tuberculosis* infection. The monoclonal antibody used (clone MP2-8F8) has been previously shown to have IL-3 neutralizing properties (1, 20). We confirmed this using an IL-3 dependent cell line (M-NFS-60) which showed a dose-dependent decrease in IL-3 dependent growth in the presence of increasing amount of anti-IL-3 antibodies in the culture medium (Figure 1A). To determine the effect of IL-3 depletion during primary mycobacterial infection, we infected C57BL/6 mice with ~100 CFU of *M. tuberculosis* (strain H37Rv) by aerosol route and injected them with either PBS, a nonbinding isotype matched antibody or IL-3 neutralizing antibody 400 μg per mouse twice a week starting from the day prior to infection. When the bacillary burdens in the lungs were compared four weeks after infection, no differences in bacillary burden among these groups were observed (Figure 1B), suggesting neutralization of IL-3 did not have any impact on bacillary control after primary infection with *M. tuberculosis*. To determine whether IL-3 depletion has an effect on BCG vaccination, the mice vaccinated with *M. bovis* BCG-Danish (2 x 10^6^ CFU by subcutaneous injection) or received sham vaccination (subcutaneous PBS injection) four weeks prior to challenge with *M. tuberculosis* (~100 CFU administered by aerosol). BCG vaccinated mice received anti-IL-3 antibody or isotype control antibody beginning on the day prior to vaccination, and subsequently, twice per week, until sacrifice, while the unvaccinated PBS control mice received injection of these antibodies only during the post-challenge period. When the bacillary burden was determined four weeks after *M. tuberculosis* challenge, a reduction in bacillary burden was observed as expected in BCG vaccinated mice. This was not significantly altered by the presence or absence of neutralizing antibodies to IL-3 (Figure 1C). Collectively, these experiments showed that neutralization of IL-3 did not have any effect on bacillary control during *M. tuberculosis* infection or that mediated by *M. bovis* BCG vaccination.

**Figure 1.**
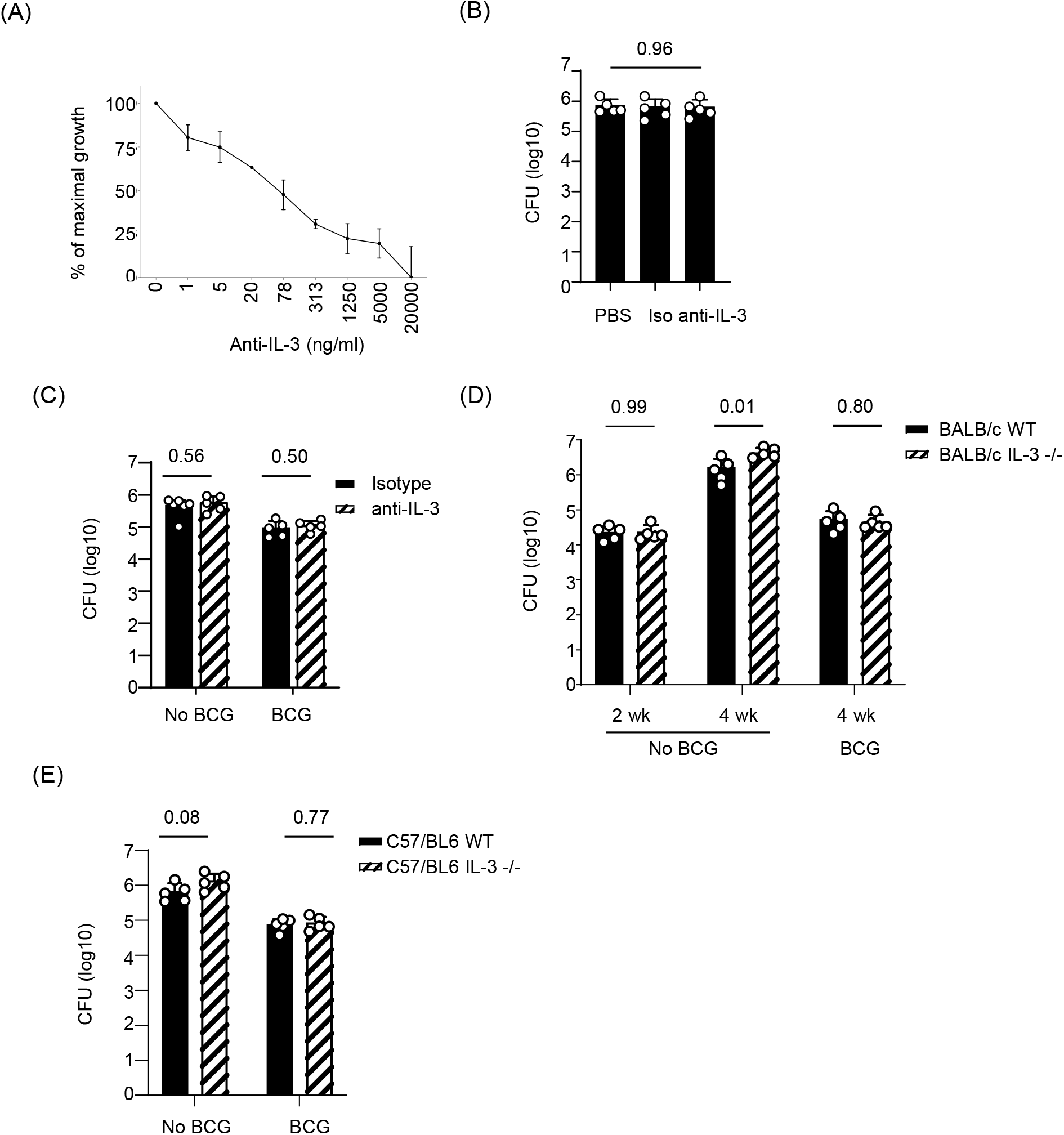
Effect of IL-3 neutralization or deficiency on *M. tuberculosis* bacillary burden. **(A)**. Neutralizing activity of anti-IL-3 antibodies (clone MP2-8F8). Varying amount of anti-IL-3 antibodies are added to M-NFS-60 cells growing in the presence of 320 U/ml of IL-3. Forty eights hours later, MTT assay is performed and the data is shown as percentage of growth obtained with the cells without anti-IL-3 antibodies. The results are representative of two independent experiments. **(B)** C57BL/6 mice received anti-IL-3 neutralizing antibody (anti-IL-3), isotype control antibody (Iso) or PBS starting from the day prior to infection. The bacillary burden in the lung at four weeks after infection is shown. The results are representative of 2 independent experiments. P value corresponds to a one-way ANOVA comparing 3 groups. **(C)** Unvaccinated or BCG-vaccinated C57BL/6 mice were challenged with *M. tuberculosis* through the aerosol route and received anti-IL-3 antibodies (anti-IL-3) or isotype control antibodies (Iso) as indicated. BCG-vaccinated mice received antibodies throughout the study period starting from the day prior to vaccination while the non-vaccinated mice received antibodies from the day prior to infection with *M. tuberculosis*. The bacillary burden in the lung at four weeks after infection is shown. The results are representative of 2 independent experiments. P values correspond to Student’s unpaired t test. **(D)** BALB/c IL-3 −/− mice or BALB/c WT mice with or without prior vaccination with *M. bovis* BCG were infected with *M. tuberculosis* by the aerosol route. Bacillary burden in the lung at two or four weeks after infection are shown. The results are representative of 3 independent experiments. P values correspond to a Student’s unpaired t tests. **(E)** IL-3 −/− or WT mice in the C57BL/6 background with or without prior vaccination with *M. bovis* BCG were infected with *M. tuberculosis* by the aerosol route and bacillary burden in the lung at four weeks after infection is shown. The results are representative of 2 independent experiments. P values correspond Student’s non-paired t tests. CFU = Colony Forming Unit, WT = wild type, IL-3 −/− = IL-3 genetic deletion.

Since neutralization by antibody may not completely block the effect of IL-3, we conducted similar experiments in *IL-3* ^−/−^ mice in a BALB/c background (BALB/c *IL-3* ^−/−^). Four weeks after *M. tuberculosis* infection, the bacillary burden was significantly higher in the BALB/c *IL-3* ^−/−^ mice compared to the WT BALB/c mice (Figure 1D). The difference in bacillary burden was not observed at 2 weeks post-infection (Figure 1D), suggesting that adaptive immunity conferred by IL-3 producing T helper cells may have been responsible for the difference seen at 4 weeks post-infection. This would be consistent with previous observations that the protection conferred by the adaptive immunity in mouse models of tuberculosis is delayed compared to many other infection models, and becomes evident approximately 3 weeks post-infection (25). This difference, however, was not seen when mice were previously vaccinated with *M. bovis* BCG (Figure 1D). Similar results were obtained when experiments were performed with IL-3 deficient mice in a C57BL/6 background (C57BL/6 *IL-3* ^−/−^), with or without prior BCG vaccination (Figure 1E).

We also assessed whether the increased bacterial burden seen in IL-3 deficient mice compared to WT mice was associated with increased lung pathology or decreased survival after *M. tuberculosis* infection. Routine histopathological examination of the lungs harvested at four weeks post-infection showed poorly formed granulomas in both WT and BALB/c *IL-3* ^−/−^ mice. However, the area of lung involved, and the lymphocytic infiltrates seen within the granulomas, were higher in BALB/c *IL-3* ^−/−^ mice compared to WT mice (Figure 2A). Consistent with this increased pathology, BALB/c *IL-3* ^−/−^ mice showed a trend toward earlier death after infection than WT mice, although this difference did not reach statistical significance (Figure 2B). No difference in survival was seen when mice were infected by intravenous route (Figure 2B), presumably because IL-3 secreting T helper cells were induced only when the infection occurred at epithelial barriers and not through blood as we have demonstrated previously (15). Collectively, these experiments suggested that IL-3 made a relatively small but detectable contribution to protection against infection with *M. tuberculosis* in standard mouse models of this infection.

**Figure 2.**
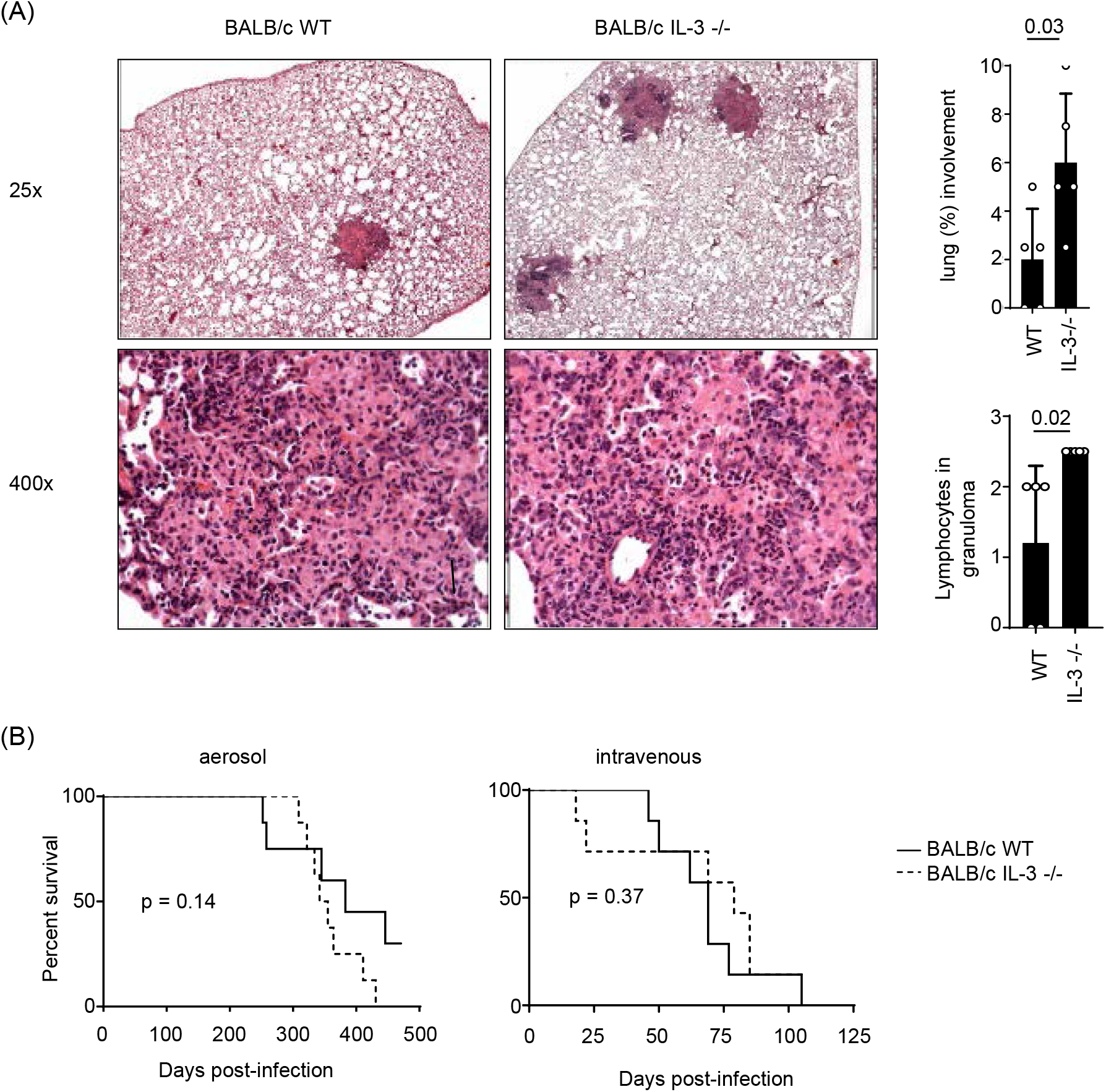
Lung pathology and survival in BALB/c IL-3 −/− mice after *M. tuberculosis* infection. **(A)**. BALB/c IL-3 −/− mice and BALB/c WT mice were infected with *M. tuberculosis* by the aerosol route and the lungs underwent routine histopathological examination at 4 weeks after infection. The micrographs represent H&E stained sections of the lung under low (25x) and high (400x) power. High power images show representative granulomas. The bar charts represents the quantification of area of involved by granulomas (above) and the percent of lymphocytes within the granulomas (below). Refer to methods for details. P values correspond to Student’s unpaired t tests. **(B).** BALB/c IL-3 −/− and BALB/c WT mice were infected by aerosol (left) or intravenous (right) route and were monitored until death. The experiment is performed once with a total of 16 (aerosol) and 14 mice (intravenous). P values correspond to Log-rank tests.

### 3.2 Role of IL-3 against infection with HSV

A protective effect of IL-3 in a subcutaneous inoculation model of HSV-1 has been shown previously using neutralizing antibodies to IL-3 (11). We first validated that observation using a modified skin scarification model in which the flank skin of mice was depilated and scarified prior to direct inoculation with HSV-1 (strain B^3^ × 1.1) using a dose of one LD90 (1 x 10^5 PFU/mouse). While all BALB/c *IL-3* ^−/−^ mice succumbed by day 7 after infection, 20% of WT mice survived to at least day 14 when the experiment was terminated (Figure 3A). Consistent with earlier death in IL-3 deficient mice, the signs of disease were more pronounced in BALB/c *IL-3* ^−/−^ mice compared to WT mice (Figure 3B). These results confirmed the protective nature of IL-3 against skin infection by HSV-1, consistent with the previous report (11).

**Figure 3.**
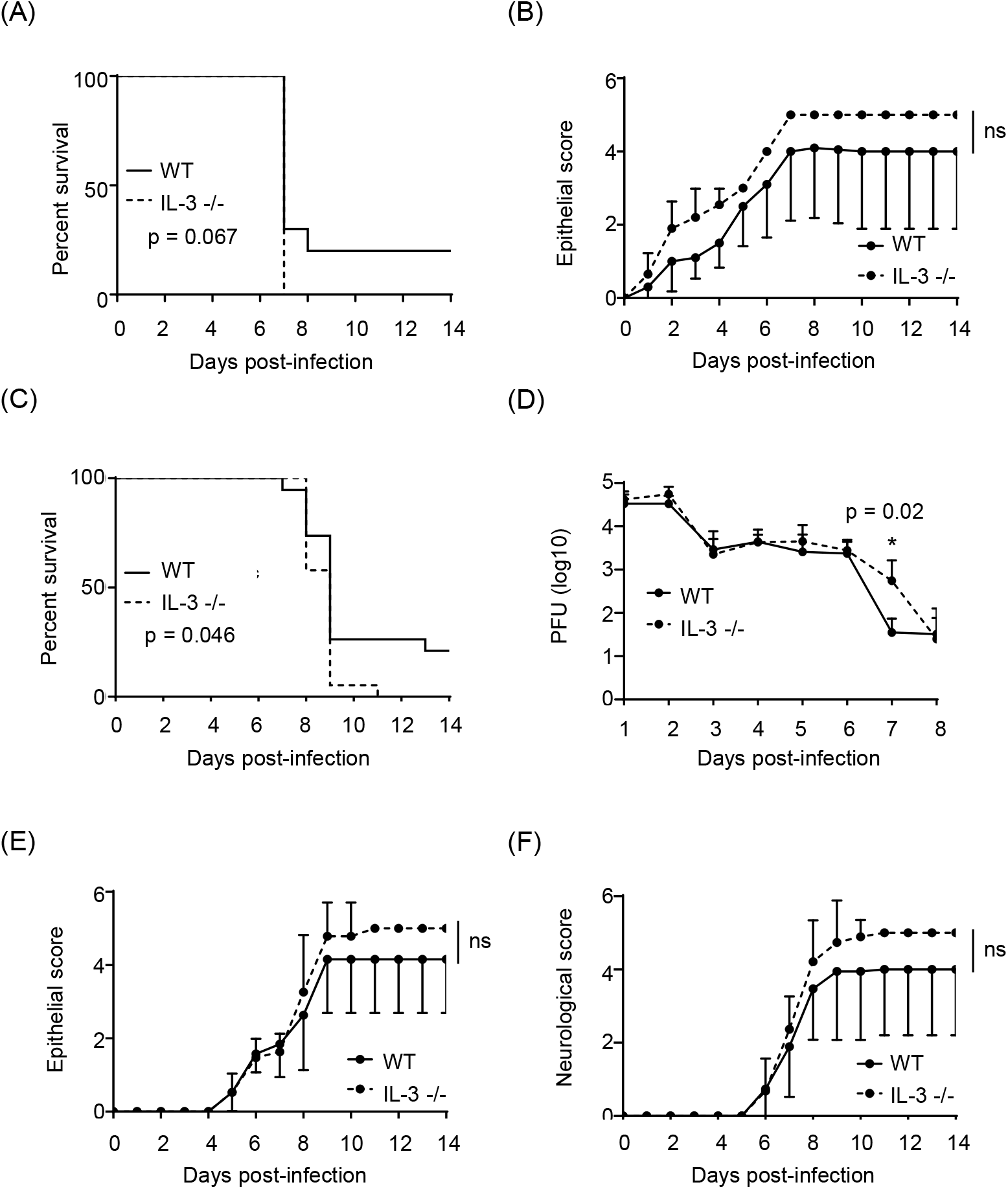
Disease burden in BALB/c IL-3 −/− mice after infection with HSV. **(A).** BALB/c WT and BALB/c IL-3 −/− male mice were infected with HSV-1 by a modified skin scarification method and monitored daily for disease development. The survival of mice are shown. The experiment is performed once with a total of 20 mice. P value represents Log-rank test. **(B).** The mean and standard deviation of the epithelial disease scores of mice in panel A are shown. The statistical significance is calculated by performing multiple t tests followed by multiple testing correction by Benjamini, Krieger, & Yekutieli False discovery rate (FDR)-approach. **(C).** BALB/c WT and BALB/c IL-3 −/− female mice were infected with HSV-2 (strain 4674) intra-vaginally and monitored daily for disease development. The survival of mice are shown. The plots show cumulative data from 2 independent experiments representing a total number of 38 nice. P value represents Log-rank test. **(D)** The viral burden of mice in panel C as enumerated from daily vaginal washes. P value indicates Students unpaired t test comparing the viral burden of WT and IL-3 −/− mice on day 7. **(E).** The mean and standard deviation of the epithelial disease scores of mice in panel C. The statistical significance is calculated as in panel B. **(F)** The mean and standard deviation of the neurological disease scores of mice in panel C. The statistical significance is calculated as in panel B.

Since the importance of IL-3 in an intra-vaginal infection model has not been reported to our knowledge, we examined this by infecting WT and BALB/c *IL-3* ^−/−^ mice at this site with a dose of one LD90 (5 x 10^4 PFU) of HSV-2 (strain 4674) and monitored for signs of disease. While all BALB/c *IL-3* ^−/−^ mice succumbed to infection by day 11 after infection, 20% of WT mice survived until day 14 (Figure 3C). Infectious virus titers from the genital tract were comparable between BALB/c *IL-3* ^−/−^ and WT mice except for the day 7, on which the burden was higher in the IL-3 deficient mice (Figure 3D). Consistent with earlier death in IL-3 deficient mice, epithelial disease and neurological disease were more pronounced in BALB/c *IL-3* ^−/−^ mice compared to WT mice (Figure 3E & 3F). These results showed a protective effect of IL-3 in an intra-vaginal infection model with HSV-2.

## 4 Discussion

In the current study, we provide evidence that in mouse models of aerosol infection with *M. tuberculosis*, IL-3 is required for optimal expression of protective immunity. Compared to WT mice, IL-3 deficient mice showed increases in both bacillary burden and lung pathology, and decreased survival after *M. tuberculosis* challenge. These differences achieved statistical significance, although the magnitude of the difference was small. This may explain why neutralization with anti-IL-3 antibodies did not yield results similar to those obtained with *IL-3* ^−/−^ mice, as the protective effect of IL-3 may be retained even with very low levels of activity that cannot be completely eliminated by antibody neutralization. We also recognize that other factors may be responsible for the differences observed between the antibody neutralization and the genetic deletion model; antibody neutralization eliminates the effects of IL-3 in a normal adult mouse for a brief period, while the genetic deletion of IL-3 may have direct and indirect effects on the immune system starting from embryonic development (26). We also noticed that the differences between *IL-3* ^−/−^ mice and WT mice were not observed if mice were previously vaccinated with BCG. While the mechanism has not been investigated, this result suggested that BCG vaccination masks the protection conferred by IL-3. Given the small magnitude of protection conferred by IL-3, we did not perform additional experiments to demonstrate directly whether IL-3 secreted by CD4^+^ T cells was responsible for the phenotype seen in the IL-3 −/− mice. Our results, however, indirectly support the hypothesis that CD4^+^ T cells are responsible for the phenotype. For example, the protective effect of IL-3 was seen at 4 weeks but not at 2 weeks after infection. If IL-3 produced by innate immune cells was responsible for the phenotype, the difference would most likely have been seen also at 2 weeks after infection since bacterial control due to adaptive immune responses occur approximately 3 weeks after infection in unvaccinated mice (25). In addition, no differences in survival were seen between *IL-3* ^−/−^ and WT mice after intravenous infections. If IL-3 from innate immune cells were protective against *M. tuberculosis*, increased mortality in *IL-3 −/−* mice compared to WT mice would be expected after intravenous infection since IL-3 secreting T helper cells were not generated in either mice (15). However, we acknowledge that different innate immune cells might be involved in the bacterial control when different routes of infections are used (27).

We also showed that IL-3 was protective in skin and vaginal infections by HSV. In the vaginal infection model using HSV-2, BALB/c *IL-3* ^−/−^ mice showed reduced survival, and increased viral burden and disease scores as compared to WT mice. As in the mycobacterial infection model, the differences were statistically significant, but the magnitude of the effect was modest. Although we have not performed additional experiments to demonstrate the involvement of IL-3-secreting CD4^+^ T cells in this phenotype, we propose that the increased viral burden seen on day 7 after infection could be caused by the absence of IL-3 secreting CD4^+^ T cells. Consistent with this, we have shown previously that IL-3^+^ T helper cells are present on day 7 after infection (15). In the modified skin scarification model using HSV-1, BALB/c *IL-3* ^−/−^ mice showed reduced survival and increased disease scores compared to WT mice. These results were consistent with previously published results in which mice which received neutralizing antibodies to IL-3 showed reduced survival compared to mice that received control isotype-matched antibodies (11). Even in different experimental conditions such as in CBA mice as previously reported, different doses (LD90 vs 5xLD50) and route of inoculation (modified skin scarification vs subcutaneous injection into foot pad) and method of IL-3 blocking (germline deletion versus neutralizing antibody), our results consistently showed that IL-3 conferred protection.

Multiple reasons may explain the low magnitude of protective effect seen with IL-3 in our models. One possible explanation is that the frequency of IL-3-producing CD4^+^ T cells is too low to provide large differences. In mycobacterial infections, only a small fraction of antigen-specific T helper cells produces IL-3 (16). This explanation, however, may not apply in HSV infections where the numbers of IFNγ and IL-3-producing CD4^+^ T cells were comparable (16). Another explanation is that the activity of IL-3 might be compensated by other cytokines, especially GM-CSF which functions similar to IL-3 and is co-secreted by IL-3 producing CD4^+^ T cells as we and others have shown (15). Other cytokines such as IFNγ, IL-2 and TNF may also partially compensate for IL-3, as most IL-3-secreting T helper cells co-secrete these cytokines (15). The questions of whether IL-3 derived from CD4^+^ T helper cells has an effect on antimicrobial immunity, or whether IL-3 represents a marker for a distinct subset of barrier-associated T helper cells, remain unresolved. However, our results support the conclusion that IL-3 is required for complete expression of protective immunity against *M. tuberculosis* and HSV infections in mouse models of these infections.

## 6 Author Contributions

SKV and SAP designed the research. SKV, TWN, NKS, MFG, PA, JX and JK performed the research. SKV and SAP analyzed the data. BCH, JC, and WRJ consulted and advised on the research. SKV, TWN, BCH, JC, WRJ and SAP wrote and reviewed the manuscript. All authors read and approved the final manuscript.

## 8 Acknowledgments

We thank Booki Min (Cleveland Clinic, OH) for kindly providing IL-3 −/− mice. We thank Fred D. Finkelman (University of Cincinnati, OH) for kindly providing MP2-8F8 hybridoma cells. We thank Rani Sellers and Amanda Beck (Albert Einstein College of Medicine, NY) for help with histopathological studies.

## 9 Supplementary Material

No supplementary material is included.

